# Glucose levels are associated with mood, but the association is mediated by ratings of metabolic state

**DOI:** 10.1101/2025.03.28.645871

**Authors:** Kristin Kaduk, Marie Kaeber, Anne Kühnel, María Berjano Torrado, Melina Grahlow, Birgit Derntl, Nils B. Kroemer

**Affiliations:** Department of Psychiatry and Psychotherapy, Tübingen Center for Mental Health, University of Tübingen, Tübingen, Germany; Section of Medical Psychology, Department of Psychiatry and Psychotherapy, Faculty of Medicine, University of Bonn, Bonn, Germany; German Center for Mental Health (DZPG), partner site Tübingen; German Center for Diabetes Research (DZD), Neuherberg, Germany

**Keywords:** continuous glucose monitoring, interoception, ecological momentary assessment, longitudinal

## Abstract

Hunger is commonly linked to negative mood and mood shifts are thought to arise from sensing the body’s internal state. However, it is unclear whether circulating glucose levels affect mood subconsciously or via subjectively sensed metabolic states. Here, we continuously monitored the interstitial glucose levels of 90 healthy adults for four weeks while they completed ecological momentary assessments (EMA; M=47 runs per participant) to rate their mood and metabolic states. As expected, hungry participants reported lower mood, and metabolic state ratings were associated with glucose levels. Although glucose levels were associated with mood, the metabolic state ratings fully mediated this association. Individual differences reflecting metabolic health (i.e., BMI and insulin resistance) did not affect the interaction between glucose and metabolic state ratings on mood. Notably, individuals with higher interoceptive accuracy had fewer fluctuations in mood ratings. We conclude that hunger-related mood shifts depend on conscious sensing of the body’s internal state instead of acting subconsciously. Our study highlights the relevance of considering the self-report of bodily signals in understanding mood shifts, offering new fundamental insights into mood regulation mechanisms.

**Key messages:** *What is already known on this topic:*

- Although glucose is linked to satiety and hunger to negative mood, it is not known if this is due to the conscious perception of metabolic states.

*What this study adds:*

- The longitudinal study design leveraging continuous glucose monitoring (CGM) and ecological momentary assessment (EMA) over four weeks, offers a unique perspective on glucose-mood interactions in naturalistic settings.
- We found that the link between mood and glucose levels is fully mediated by consciously sensed metabolic states, clarifying the interoceptive link between glucose and mood.

*How this study might affect research, practice or policy:*

- Mood is shaped by perceived metabolic states highlighting the relevance of metabolic health for mental health.
- These insights improve our understanding of the association between metabolic and mental disorders.

**Graphical abstract:** 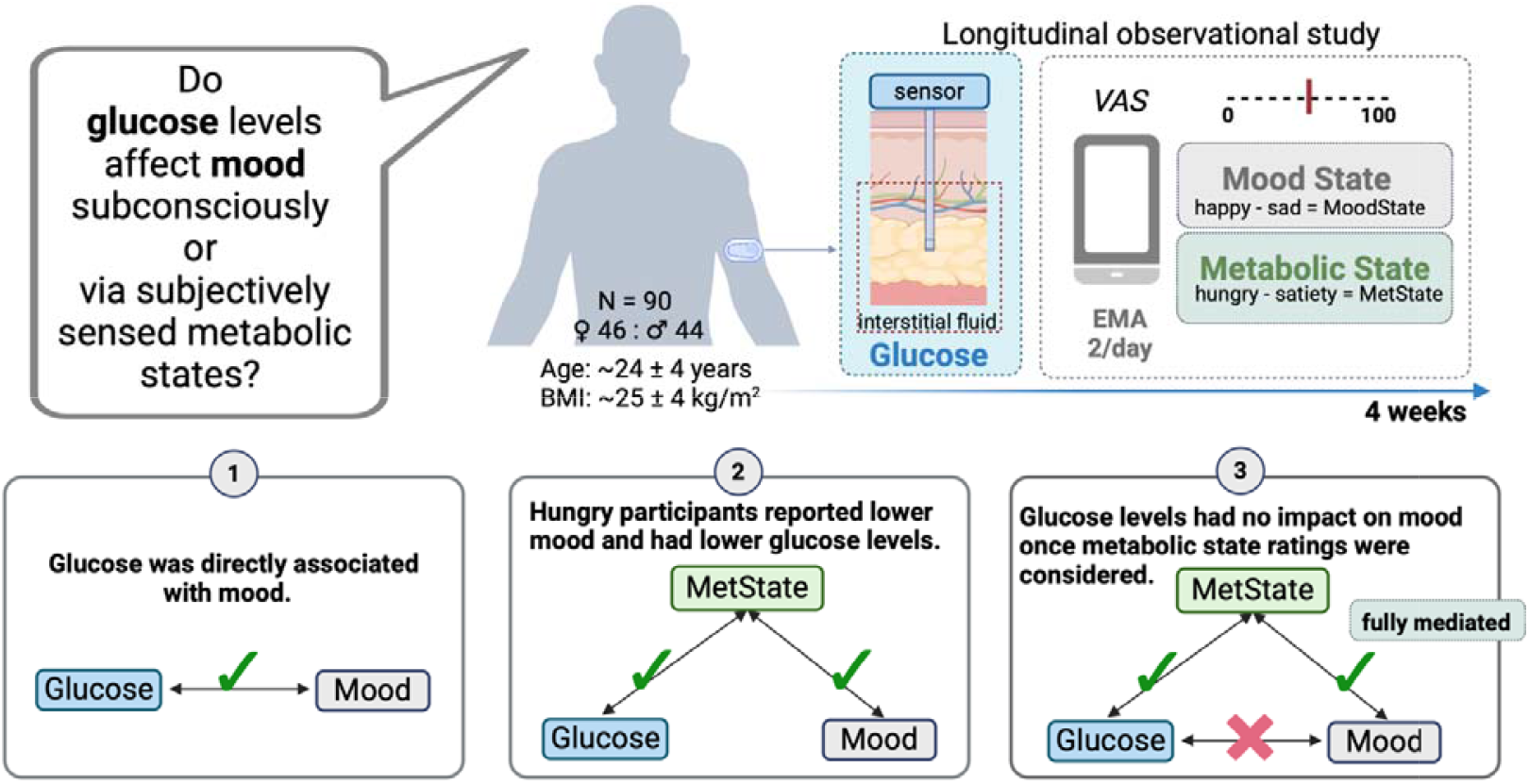

## Introduction

Hunger often precipitates mood changes, likely serving a vital function (MacCormack & Lindquist, 2019; Piccolo et al., 2020) by signaling the body’s energy demands and guiding food intake (Bennett et al., 2022; Eldar et al., 2016; Teckentrup & Kroemer, 2024). Energy metabolism is crucial for mental health, with metabolic disorders linked to psychological sequelae (Lyra E Silva et al., 2019; Vancampfort et al., 2015). Central to this interaction between energy metabolism and mood is glucose, the body’s primary energy source (Dienel, 2019; Mergenthaler et al., 2013), influencing mood (Gold et al., 1995; mood under cognitive demands: Owen et al., 2012; Owens, 1997), aggression (Bushman et al., 2014; Denson et al., 2010), and tension (Benton & Owens, 1993; Gold et al., 1995). Despite the importance of body-brain interactions for mental health and well-being, whether circulating glucose affects mood beyond self-reported metabolic states remains unclear.

The homeostatic view states that physiological signals, such as glucose, shape hunger and mood to counteract a negative energy balance. Low glucose levels trigger hormone release (e.g., glucagon) to mobilize glucose from liver stores, inhibit insulin secretion, stimulate appetite, and facilitate eating in rats (Campfield et al., 1985; Campfield & Smith, 1986) and humans (Melanson et al., 1999; Schultes et al., 2003). Elevated glucose levels stimulate insulin secretion inducing satiety, decreasing tension (Benton & Owens, 1993), and appetite (Woods et al., 2006). However, laboratory studies on mood-related effects of glucose (Van De Rest et al., 2018), often using high-calorie drinks or food, have yielded inconclusive results (Benton & Owens, 1993; Green et al., 2001; Markus, 2007, 2007; Owen et al., 2012; Owens, 1997; Scholey et al., 2014; Seo et al., 2014; Ullrich et al., 2015). Continuous glucose monitoring (CGM) enables research beyond controlled, but artificial settings (Keshet et al., 2023; Shilo et al., 2024). Combining CGM with ecological momentary assessment (EMA) offers a novel approach, facilitating naturalistic studies of daily mood fluctuations while reducing recall biases. For example, in patients with type 2 diabetes, higher glucose levels were associated with negative mood (Wagner et al., 2017). In contrast, healthy adolescents showed elevated mood and reduced fatigue with higher glucose levels (Zink et al., 2020). These findings indicate a link between mood and glucose levels but the role of sensed hunger is not considered.

Hunger motivates energy intake in response to ingestion-related internal cues (Aunger & Curtis, 2013; Beaulieu & Blundell, 2021), acting as both a homeostatic signal and an allostatic signal to anticipate negative excursions in energy balance (e.g., low energy supply). In animals, metabolic states influence behavior and brain responses. Agouti-related peptide (AgRP) neurons in the hypothalamus signal hunger’s negative valence (Betley et al., 2015). Energy intake restores glucose levels, modulating the activity of metabolic-sensing AgRP neurons and activating the dopaminergic mesolimbic system (Reichenbach et al., 2022), shifting hunger’s negative valence state into positive reinforcement. In humans, hunger and satiety are intricately linked to mood (Monello & Mayer, 1967; Swami et al., 2022). Hunger is related to restlessness, excitability (Monello & Mayer, 1967), sadness (De Rivaz et al., 2022), increased irritability, aggression, and anger (Nettle, 2017; Swami et al., 2022), whereas satiety promotes satisfaction, relaxation, and calmness (Monello & Mayer, 1967). Accordingly, hungry women with healthy weight reported higher tension, anger, fatigue, and confusion and lower vigor and esteem (Ackermans et al., 2022).

Findings suggesting that hunger contributes to negative mood are consistent with the psychological construction of emotion theory (see Barrett, 2006, 2016). It posits that homeostatic processes (e.g., hunger as a signal for energy imbalance) generates core affect, shaping mood through bodily sensations and external stimuli (MacCormack & Lindquist, 2017, 2019; Russell, 2003). For example, MacCormack & Lindquist (2019) found that hunger biases perception and judgment, heightening negativity, particularly when individuals were unaware of their emotions showing that interoception is crucial (Craig, 2002; Feldman, Bliss-Moreau, et al., 2024; Feldman, Ma, et al., 2024). Instead, the psychological construction of emotion theory posits that interoception transforms physiological states (e.g., low blood sugar) into the cognitive experience of hunger by integrating bodily sensations with contextual information (Barrett & Simmons, 2015; MacCormack & Lindquist, 2017; Simmons & DeVille, 2017). Interoceptive accuracy in detecting metabolic changes therefore likely influences mood regulation as it arises from sensing bodily signals.

To summarize, glucose surges are associated with satiety and hunger is linked to negative mood states. However, it remains unclear whether conscious processes reflected in subjective ratings of metabolic state fully mediate the association between glucose and mood. To address this gap, we conducted a four-week study combining CGM and EMA in naturalistic settings to examine the association of glucose levels and metabolic state ratings on mood (happiness and sadness). Consistent with previous findings, we expected lower glucose levels to be associated with decreased metabolic state ratings and mood. Our study shows that the association of glucose with mood was fully mediated by metabolic state ratings. Moreover, higher interoceptive accuracy was associated with fewer mood fluctuations, highlighting the role of interoception in mood regulation in accordance with the psychological construction of emotion theory.

## Methods

We preregistered our study protocol at the Open Science Framework (https://osf.io/gpr52). The reported data are part of a larger study examining the potential link between metabolic states and dopamine-related reward learning, and the reported analyses were not preregistered.

### Participants

The sample size was selected to provide at least a power of 1-β = .94 to study small-to-moderately sized within-subject effects (*d*_z_~.40), leading to a lower-bound estimate of 80 participants after quality control. A total of 97 participants were invited to participate. Participants were included in the analysis if we collected at least 20 EMA runs with concurrent CGM data. Consequently, we excluded 7 participants due to extended malfunction of CGM (e.g., detached sensors or loss of transmission). This led to a sample of 90 participants (46 women, *M*_age_: 24.27 ± 3.57 years, range: 18–34, *M*_BMI_: 24.71 ± 4.09 kg/m^2^, range: 18-36.7). According to screenings, all participants were physically and mentally healthy (except for a heightened BMI) and reported no history of neurological, neurosurgical, or cardiological disease or treatment. Participants received fixed compensation of 160€ or partial course credits and additionally performance-dependent wins from the tasks and EMA. All participants provided their informed written consent before the experiment. The ethics committee of the Faculty of Medicine at the University of Tübingen approved the experiment, and all procedures were carried out in accordance with the Declaration of Helsinki.

### Experimental procedure

Participants had five laboratory sessions over four weeks (Fig. 1), each occurring once a week, followed by two MR sessions. In the first session (S1), they completed three tasks (Effort Allocation Task (Neuser et al., 2020), Go/Nogo Reinforcement learning paradigm (Kühnel et al., 2020), Reward Ratings Task (Müller et al., 2022), and state questions. Participants were fitted with a CGM sensor during the first session (FreeStyle Libre 3 sensors, Abbott GmbH, Abbott Diabetes Care, Wiesbaden, Germany). They received a cell phone to track their food intake, synchronize the data of the sensor via the FreeStyle Libre app and play Influenca. EMA questions regarding mood and metabolic states were included in the gamified reinforcement learning task, ‘Influenca’ that they were asked to play twice a day over the following four weeks with a minimum gap of 2 h between assessments (Neuser et al., 2023). Before each run of ‘Influenca’, participants completed several questions capturing momentary metabolic (hunger, satiety, thirst, time since the last meal, and consumption of coffee or snacks in the previous two hours), mental (mood including self-satisfaction, happiness, sadness, stress), and physical (physical health) states at that moment. Participants indicated their response on a visual analog scale, ranging from 0, representing “not at all”, to 100, representing “very much”. During the third session (i.e., after the second week), the first glucose sensor was replaced and then removed after two weeks. If a sensor fell off or was no longer reporting data, participants scheduled an appointment for a replacement. To determine fasting glucose, insulin, and triglyceride levels, we drew blood following a 12 h overnight fast at three different visits.

**Figure 1.**
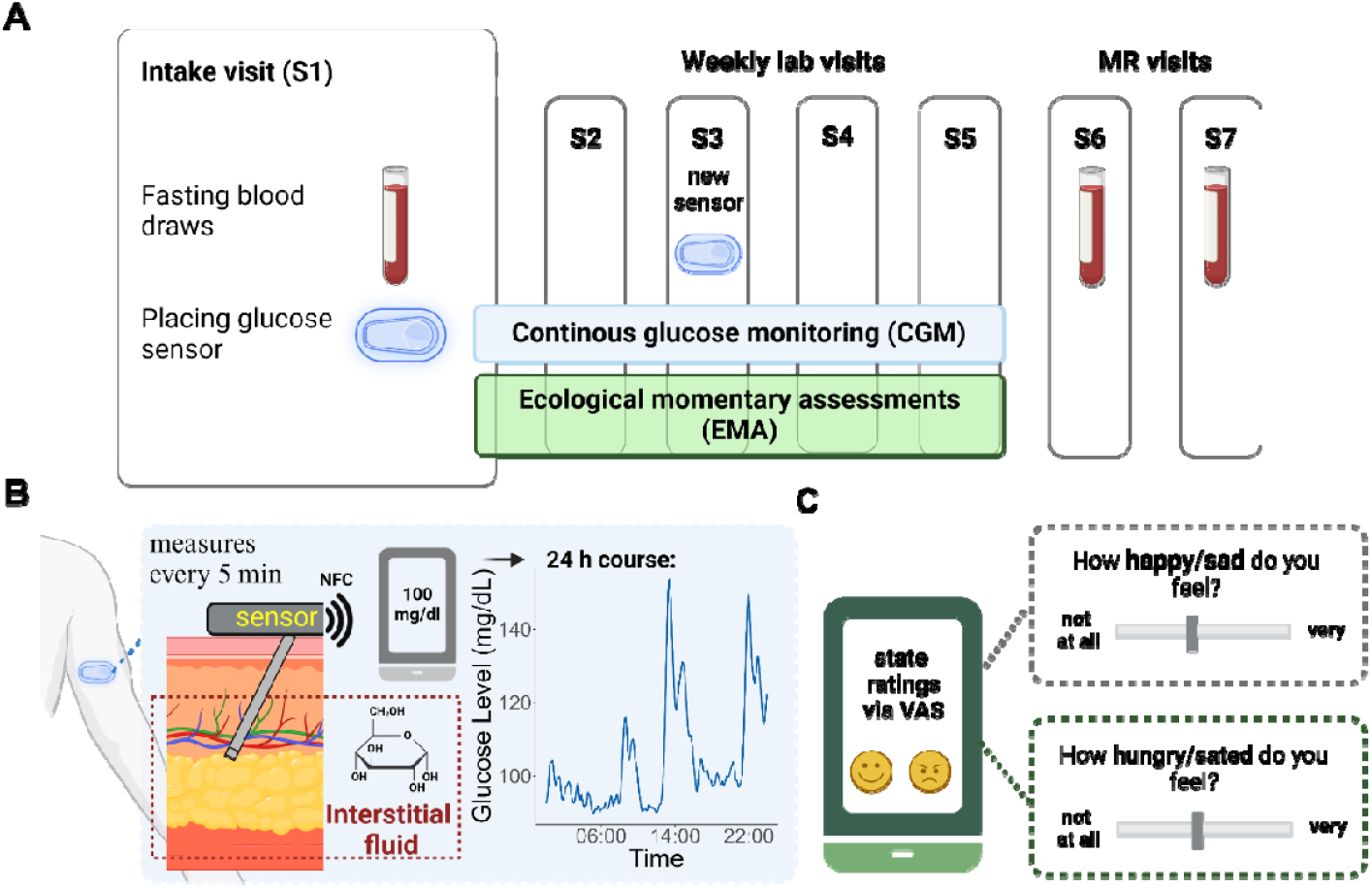
Summary of the study. **A:** The schematic shows the experimental procedure, illustrating five weekly lab visits followed by 2 MR visits labeled from S1 to S7. After the first session (S1), participants undergo four weeks of continuous glucose monitoring (CGM) combined with ecological momentary assessment (EMA). **B:** The CGM system uses a small flex sensor placed at the posterior upper arm that is inserted in the subcutaneous fatty tissue. The glucose sensor, positioned beneath the skin surface layers (epidermis and dermis), measures glucose levels in the interstitial fluid within the subcutaneous adipose tissue. The data is wirelessly sent via near field communication (NFC) to the FreeStyle Libre 3 app on the participant’s cell phone. The accuracy of the sensor concerning capillary glucose has been validated (Alva et al., 2022). **C:** Participants rate their metabolic and mood states twice daily using a visual analog scale (VAS), ranging from 0 (not at all) to 100 (very). Created with BioRender.com.

## Data analysis

### Preprocessing of the glucose data

We extracted raw sensor glucose values according to best-practice recommendations (Visser & Gillard, 2024). To account for the variability from different sensors and missing data, we first segmented the data into sections marked by intervals longer than 20 min. To rectify and normalize the glucose recordings and mitigate spurious fluctuations, we used MATLAB’s ‘msbackadj’ function from the Bioinformatics toolbox. To adjust each segment independently to correct for shifts, the ‘msbackadj’ function estimates a baseline for each segment and adjusts the segment. Next, we added a stable baseline value to these corrected glucose values for each participant. To establish this stable baseline for each participant, we identified the glucose period with the lowest glucose variability using the raw glucose values during the overnight fasting window (midnight to 8 am). This was done using a sliding window, of 1 h duration and moving in steps of 5 min. After baseline correction, we implemented Gaussian smoothing with a window length of 5 data points to reduce noise while preserving the signal’s essential characteristics. Subsequently, we partitioned the adjusted data into standardized five-minute intervals for further investigation. As the physiological delay of glucose transport from the vascular to the interstitial space is 5-6 minutes in healthy adults (Basu et al., 2013), we extracted the CGM value, which was recorded 5-10 minutes after the corresponding EMA.

### Behavioral data

To calculate a composite mood state, we used the ratings of the “happy” item and the “sad” item to calculate for each time point the “mood state” = (happy – sad)/100. Likewise, we calculated “metabolic state” = (hunger – satiety)/100. To explore the extent to which physiological changes in glucose levels are accurately perceived, we computed the interoceptive accuracy for each individual. This was done by extracting the individual estimates of the slope predicting the metabolic state from a linear mixed-effects model of *z*-standardized glucose levels (slopes were inverted so that higher scores reflect better accuracy similar to interoceptive coherence in Young et al., 2021).

### Insulin resistance derived from blood

As a measure of insulin resistance (i.e., interindividual differences in how well the body utilized glucose), we calculated the homeostasis model assessment of insulin resistance (HOMA-IR; higher HOMA-IR reflects lower insulin sensitivity) using the glucose and insulin levels measured from fasting blood samples (Matthews et al., 1985). Most participants had three fasting blood samples to compute the median HOMA-IR, though one participant had only two samples, and nine subjects had only one. Since the distribution of the HOMA-IR and glucose levels were skewed, we ln-transformed them for parametric analyses (Emoto et al., 1999; Kroemer et al., 2013). To isolate insulin resistance from other correlated covariates, we residualized HOMA-IR values by adjusting for BMI, sex, and age.

### Statistical analyses and software

We preprocessed CGM data with MATLAB vR2022b. All statistical analyses were conducted with R (v4.3.1, R Core Team, 2023). To partition the variance into between-person and within-person effects, we computed linear mixed-effects models using lmerTest (Kuznetsova et al., 2017). To facilitate direct comparison across different scales and variables, we *z*-standardized predictors across all observations. We included random intercepts and slopes for glucose levels and metabolic state to account for the inter-individual variance in repeated measurements. All LMEs included BMI, sex, and age as covariates. To evaluate whether glucose levels alone are associated with mood, we modeled mood state as a function of glucose, followed by analyzing happy and sad separately to capture differential glucose effect on positive and negative mood. Next, we analyzed the indirect pathways of mediation by examining how glucose levels influence hunger, satiety, and metabolic state ratings, and how metabolic state ratings affect mood states using LMEs. We tested whether metabolic state ratings mediate the association between glucose levels and mood and examined insulin resistance as a moderator by including HOMA-IR values and its interaction with glucose and metabolic state. To evaluate whether interoceptive accuracy (i.e., the correspondence of changes in glucose and hunger) is associated with mood state, we modeled average mood ratings (i.e. mean) and fluctuations in mood ratings (i.e. variability) as a function of interoceptive accuracy, including BMI, age and sex as covariates (with interactions with interoceptive accuracy). We considered α ≤ .05 as significant. Data was visualized using ggplot2 (Hadley, 2016), ggridges (Wilke, 2017), ggpointdensity (Kremer, 2019), and ggside (Landis, 2021).

## Results

To investigate the association of glucose and metabolic state with mood, participants were asked to rate their states (Fig. 1C) while we recorded glucose levels using CGM (Fig. 1B). The analyses incorporated 4299 EMA runs from 90 participants. Most EMA (on average 47.76±8.83 measurements per participant, range: 22-60) were recorded (Fig. 2A).

**Figure 2.**
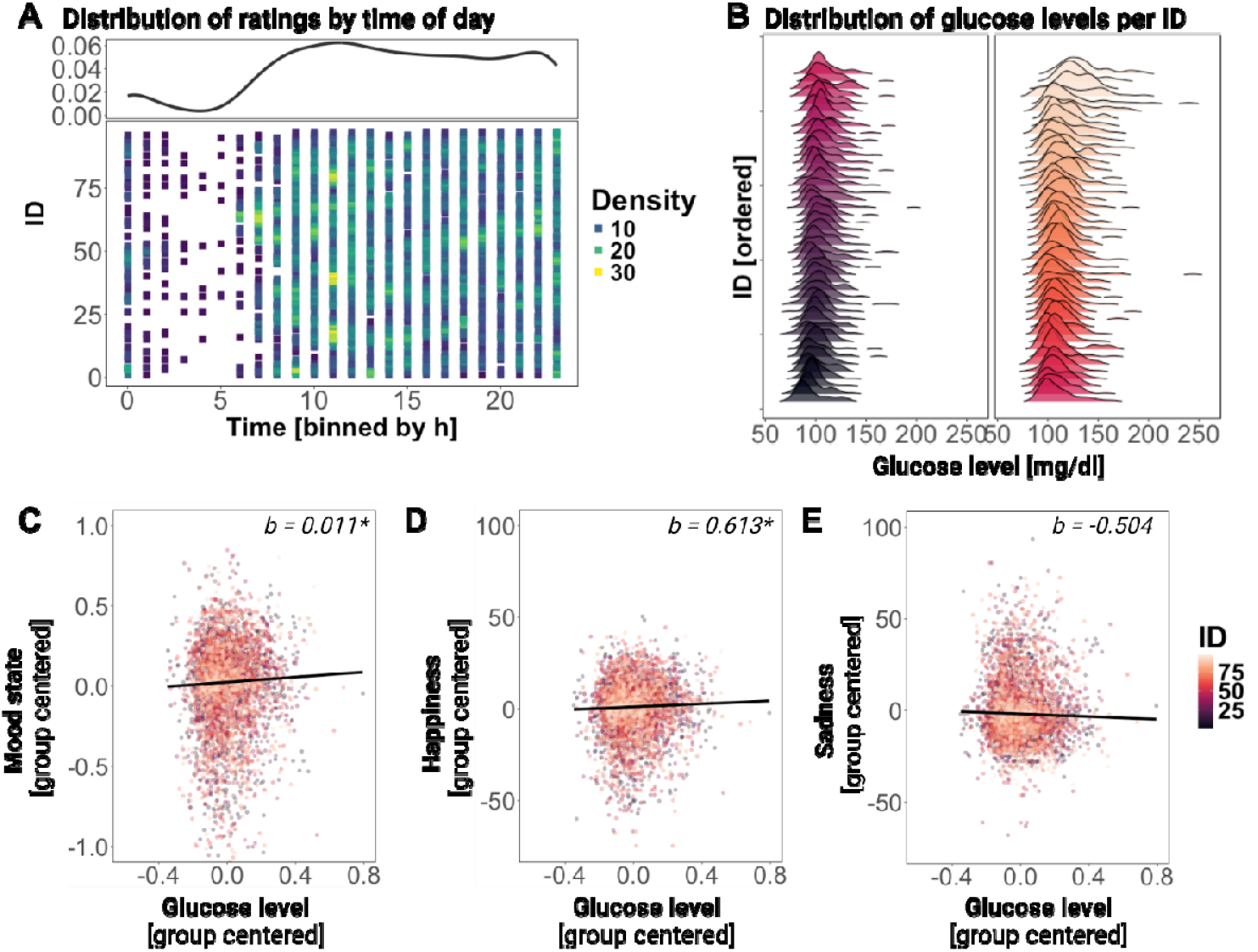
Glucose levels are associated with mood. **A:** The color of each square represents the density of measurements for each participant across the day, binned by hours. The measurements are distributed and displayed in the average density curve during the hours of usual wake time. **B:** This density plot illustrates in two rows the distribution of glucose levels corresponding to the EMA for each participant, ordered by their median glucose levels. The lower panel displays the relationship between glucose levels and **C:** mood state, **D:** happiness, and **E:** sadness, indicating that when glucose levels are high, participants reported higher mood. Each dot depicts an observation with a color-coding of participants. For visualization, mood state, happiness, and sadness are group centered, and glucose levels are ln-transformed and group-centered to match the linear mixed-effects model. Black lines indicate the relationship (fixed effect) across participants, indicating that with higher glucose levels participants are happier. * *p* < .05, ** *p* < .01.

### Mood improves with higher glucose levels

To evaluate whether glucose levels are associated with mood, we initially examined the direct effect of glucose levels on mood as outcome with a linear mixed-effects model (including BMI, age, sex, and HOMA-IR and their interactions with glucose levels to account for individual differences; *p*s > .05). Higher glucose levels were related to a better mood state (*b* = 0.011, *t*(79) = 2.08, *p* = .040, Fig. 2C; higher scores indicate more happiness vs. sadness). To explore whether there is a specific association with positive or negative mood, we estimated separate models and observed slightly stronger associations with happiness (*b* = 0.613, *t*(75) = 2.36, *p* = .021, Fig. 3D), whereas there was no significant association with sadness (*b* = −0.504, *t*(77) = −1.43, *p* = .157, Fig. 3E). To examine potential non-linearity of the data, we used a Generalized Additive Model (GAM) which confirmed that the relationship between glucose and mood is adequately described by a linear model (see Fig. S1 in the Suppl. Material).

**Figure 3.**
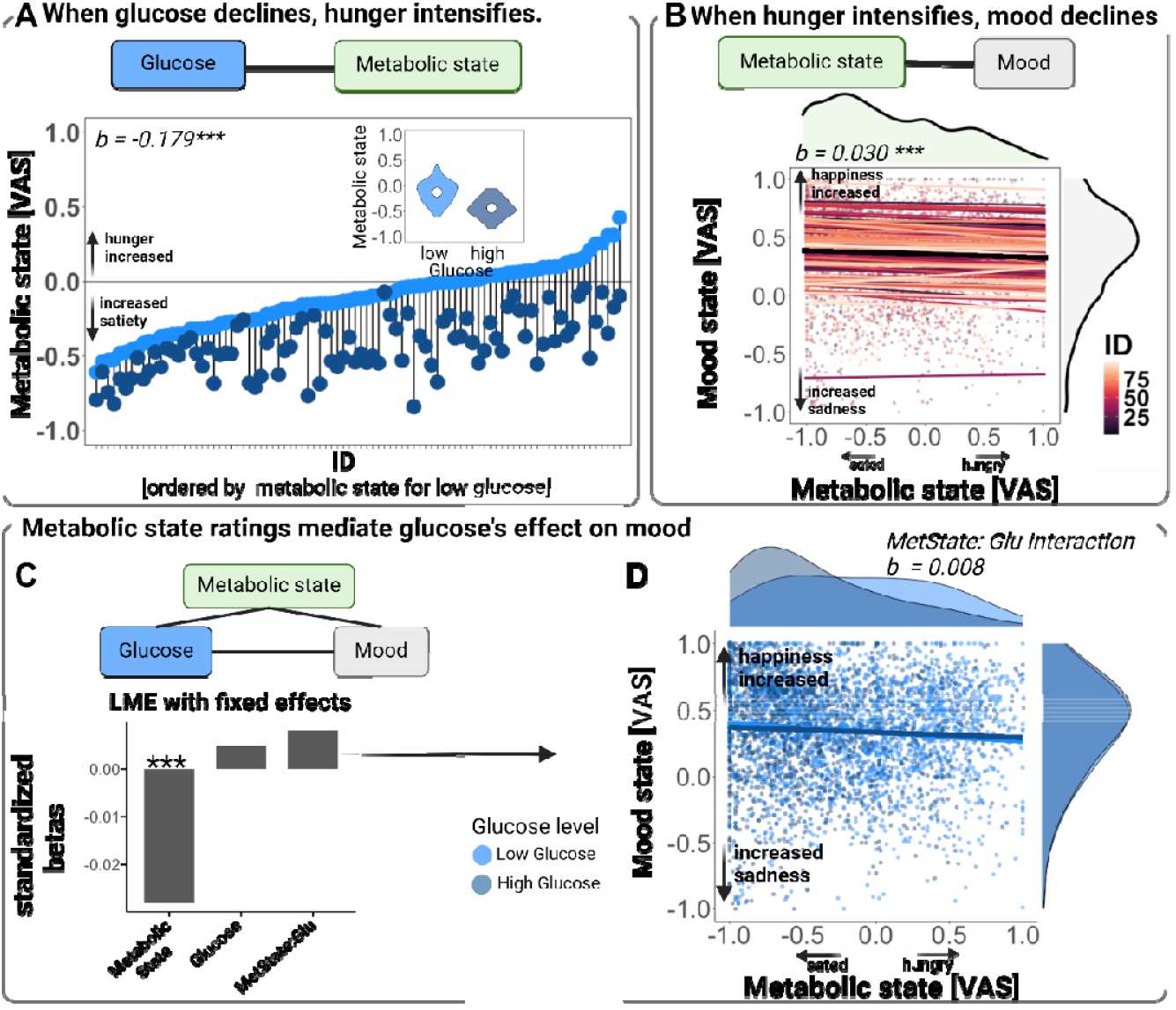
Metabolic state and glucose levels, but not their interaction, are associated with lower mood. **A:** The spaghetti plot displays the significant relationship between metabolic state ratings and mood state. Each dot depicts an observation with a color-coded grouping of participants. Colored lines show participants-specific estimates (random effect), while black indicates the average relationship (fixed effect) across participants. On top and right is the density illustrated by the distribution of metabolic and mood states, respectively. **B:** Presents the metabolic states of each participant for the low and high glucose level, which are ordered according to the participant’s state score for their low glucose level. Dots represent mean state values for the participant-specific low glucose (light blue) or high glucose (dark blue) level. To visualize differences in average metabolic state ratings between high and low glucose values, participants were divided into two equally sized groups based on their group-centered glucose values (low vs. high glucose level). **C:** Standardized linear mixed-effects coefficients of metabolic state, ln-transformed glucose, and their interaction. **D:** The combination of scatter and density plots display the interaction of metabolic state rating and glucose level on mood state, where the glucose levels are separated into low (light blue) and high (dark blue) glucose levels within each participant. * p < .05, ** p < .01. Created with BioRender.com.

### Hunger is associated with glucose levels and lower mood

To evaluate how strongly glucose levels are associated with hunger, we used a linear mixed-effects model with metabolic state ratings as outcome and glucose levels as predictor. As expected, higher glucose levels were associated with less hunger (*b* = −8.449, *t*(82) = −17.23, *p* < .001) and more satiety (*b* = 9.505, *t*(74) = 16.90, *p* < .001), leading to a strong association with the metabolic state (hungry – sated, *b* = −0.178, *t*(77) = −17.84, *p* < .001, Fig. 3A). To consider inter-individual differences, an extended model also included BMI, age, sex, and HOMA-IR and their interactions with glucose levels. Males had a higher metabolic state compared to females dependent on age and BMI (BMI × Sex × Age, *b* = −0.089, *t*(70) = −2.20, *p* = .031; main effect of sex: *b* = 0.153, *t*(75) = 3.73, *p* < .001). However, none of the covariates showed a significant interaction with glucose levels (*p*s > .05).

To evaluate whether hunger and mood are associated, we analyzed the effect of metabolic state on mood (higher scores indicate more happiness vs. sadness) using a linear mixed-effects model including only metabolic state as the predictor. As expected, when participants were hungrier, they also reported lower mood (*b* = −0.030, *t*(93) = −4.83, *p* < .001, Fig. 3B). To consider inter-individual differences, an extended model also included BMI, age, sex, and adjusted values of HOMA-IR. None of the covariates did affect significantly mood state ratings (*ps* > .05). To sum up, these findings indicate that glucose levels are a physiological signature of metabolic state, and both glucose levels and metabolic state are associated with mood.

### Metabolic state ratings mediate the effect of glucose on mood

To investigate whether the association between glucose levels and mood is mediated by ratings of metabolic state, we performed a mediation analysis (the results of the indirect pathways are reported above). We applied a linear mixed-effects model, including metabolic state, glucose level, and their interaction as predictors (Fig.3C). Glucose levels alone were no longer significantly related to mood (*b* = 0.004, *t*(82) = 0.69, *p* = .494) when self-reported metabolic state was also included in the model. However, the effect of metabolic state ratings on mood (i.e., worse mood when hungrier, *b* = −0.028, *t*(97) = −4.11, *p* < .001) was still significant independent of their glucose level. The interaction between glucose levels and metabolic state did not significantly influence mood (*b* = 0.008, *t*(104) = 1.33, *p* = .188, Fig. 3D), indicating that glucose levels do are not affected more strongly when glucose levels and metabolic state rating align.

To examine whether insulin resistance modulates the link between glucose, metabolic state, and mood, we included the adjusted values of HOMA-IR and its interactions with glucose levels and metabolic state ratings, sex, age and BMI. We found no significant interaction between HOMA-IR, glucose levels, and metabolic state on mood (*b* = 0.008, *t*(116) = 1.27, *p* = .207) and no significant main effect (*ps* > .05). Taken together, our findings suggest that a conscious self-reported metabolic state fully mediates the association between glucose and mood.

### Interoceptive accuracy is associated with fluctuations in mood

Since the relationship between glucose and mood is fully mediated by the self-reported metabolic state (i.e. interoception), we explored whether interoceptive accuracy explained associations with average mood ratings (i.e. mean) and fluctuations in mood ratings (i.e. variability). First, interoceptive accuracy was computed by extracting the individual estimates from a linear mixed-effects model, where differences in z-standardized glucose levels were predicted by the z-standardized metabolic state ratings (Fig. 4A) and inverted the estimates so that higher values reflect better accuracy. To investigate potential factors influencing the predictor of interoceptive accuracy, we examined the association between interoceptive accuracy and BMI, HOMA-IR, sex, and age. As expected, interoceptive accuracy was lower with a higher BMI (*b* = −0.036, *t*(74) = −2.88, *p* = .005, Fig. 4C), but there were no associations with HOMA-IR alone (*b* = −0.013, *t*(88) = −1.02, *p* = .312, Fig. 4B), sex (*b* = 0.038, *t*(88) = 1.62, *p* = .110) or age (*b* = 0.0003, *t*(88) = 0.02, *p* = .984). However, there was a significant interaction between HOMA-IR (adjusted for BMI, age, and sex), BMI, and sex (*b* = 0.071, *t*(88) = 2.39, *p* = .019, Fig. 4D), indicating that the interoceptive accuracy was reduced for females with high BMI and low HOMA-IR and for males with high HOMA-IR. When examining mood, we found that interoceptive accuracy was not associated with average mood ratings (i.e., mean, *b* = −0.004, *t*(88) = −0.01, *p* = .991, Fig. 4B). Instead, interoceptive accuracy was associated with fluctuations in mood ratings (i.e. variability, *b* = −0.286, *t*(88) = −2.56, *p* = .012, Fig. 4B). To sum up, interoceptive accuracy seems to play a crucial role in fluctuations in mood rather than average mood state.

**Figure 4.**
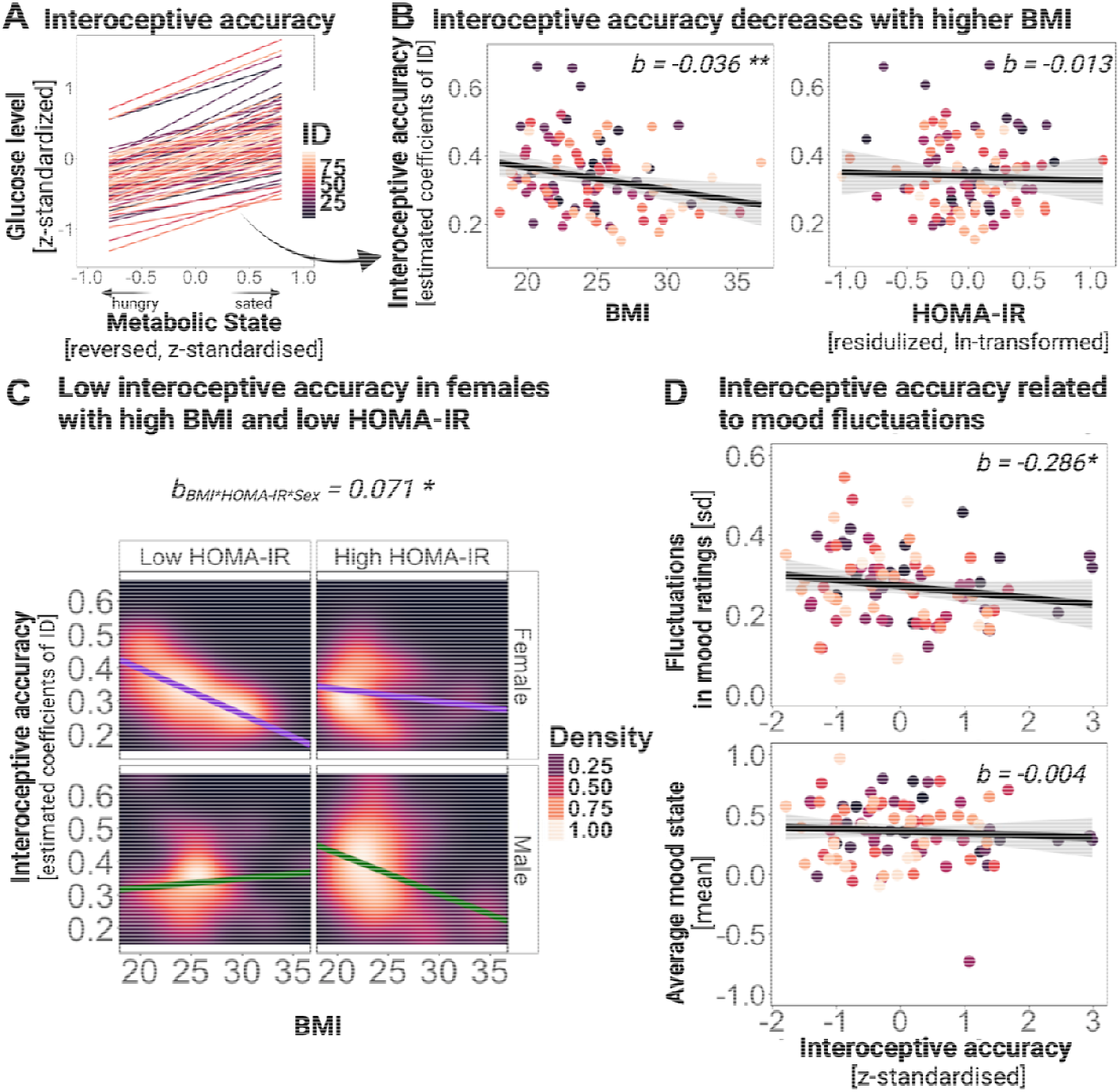
Effect of metabolic factors (BMI, HOMA-IR) on interoceptive accuracy and its association with fluctuations in mood. **A:** Depicted are the individual standardized slopes (i.e. interoceptive accuracy) as individual lines derived from a mixed-effect model, where the differences in glucose levels (ln-transformed, z-standardized) were predicted with the metabolic state ratings (reversed, z-standardized). **B:** Interoceptive accuracy is associated with BMI, but not with ln-transformed and residualized HOMA-IR. Each dot depicts an observation with a color-coded grouping of participants. The regression lines were computed via a robust linear regression method and are displayed with 95% confidence intervals. **C: I**nteroceptive accuracy is associated with BMI more strongly in females with lower HOMA-IR, indicating an interplay of metabolic health and sex on interoceptive accuracy. The plot shows the density distribution of interoceptive accuracy (on the y-axis) against BMI (on the x-axis), which is stratified across two levels of HOMA-IR by splitting participants into two equally sized groups based on their residualized HOMA-IR (low vs. high HOMA-IR. Each represents a distinct subgroup within the study population. The regression lines were computed via a robust linear regression method and are displayed with 95% confidence intervals. The color gradient of the background density ranges from black, indicating low density, to white, which indicates a high density of observations within the BMI and interoceptive accuracy. **D:** Interoceptive accuracy (z-standardized, grand mean centered) is associated with fluctuations in mood ratings (standard deviation within a person) and the average mood state (mean rating per participant).

## Discussion

Hunger worsens mood, but whether this is due to consciously sensed hunger or subconscious changes in glucose levels remains unclear. Using a deep phenotyping approach in naturalistic settings, we found an association between glucose and mood that was attenuated when metabolic state ratings were considered. This suggests that self-reported metabolic state mediates glucose’s effects on mood, refuting the idea of a largely subconscious influence. In support of an allostatic process subserving energy and mood regulation, we found that associations with positive mood were more pronounced compared to negative mood and the linear association speaks against a restricted negative valence signal as posited by homeostatic theories. We conclude that hunger-induced mood changes reflect conscious sensing of metabolic state, with hunger acting as an early metabolic signal that triggers a behavioral and affective cascade to tune adaptive behavior (Teckentrup & Kroemer, 2024)

Consistent with prior research, we observed a strong association between hunger and lower mood in healthy adults (De Rivaz et al., 2022; MacCormack & Lindquist, 2019; Swami et al., 2022). While traditionally seen as a drive to reduce energy deficits (e.g., Aunger & Curtis, 2013), hunger is now recognized as intertwined with mood (Fahed et al., 2023). Hunger ratings reflect multidimensional interoceptive sensing, integrating bodily cues (e.g., rumbling stomach, feeling cold, fatigue, irritability) (Hams & Wardle, 1987; Monello & Mayer, 1967; Stevenson et al., 2023). People process interoceptive signals across focal bodily sensations (e.g., empty stomach), diffuse sensations (e.g., nausea), and affective states (irritability, boredom) (Stevenson et al., 2015, 2023). Thus, glucose levels correlated robustly with metabolic state ratings, in line with prior research (Campfield et al., 1996; Lemmens et al., 2011; Nederkoorn et al., 2009; Schultes et al., 2003). This aligns with the psychological construction of emotion theory, which posits emotions emerge from interpreting internal states and external cues (Barrett, 2006). Accordingly, homeostatic processes (e.g., hunger signaling energy deficit) generate core affects that shape generalized mood states and the basis of emotions (Russell, 2003). This effect is pronounced when individuals lack awareness of their bodily needs (Ackermans et al., 2022; MacCormack & Lindquist, 2019). However, metabolic state ratings fully mediated the association between mood and glucose, indicating that conscious sensing of metabolic states affects mood, not glucose per se. This aligns with meta-analytic evidence refuting subconscious links between mood, self-control, and glucose depletion (Dang, 2016; Kurzban, 2010; Orquin & Kurzban, 2016; Raichle & Mintun, 2006) and challenging broad claims about direct links with self-control (Gailliot et al., 2007; Penckofer et al., 2012). Taken together, our findings support the psychological construction of emotion theory by demonstrating that hunger contributes to mood decline via conscious sensing.

Interoception plays a crucial role in mood regulation and well-being (Critchley & Garfinkel, 2017; Nayok et al., 2023; Quadt et al., 2018). Our findings show that higher interoceptive accuracy of glucose levels are associated with fewer mood fluctuations but not average mood states. This partially aligns with Young et al. (2019), who observed that individuals with higher interoceptive accuracy had reduced hunger and anxiety after glucose intake. Neither Young et al. (2019) nor Zamariola et al. (2019) found a link between interoceptive accuracy (via heartbeat counting task) and mood, further supporting our finding that interoceptive accuracy is not associated with average mood state. While most studies focus on cardiac interoception (Barrett et al., 2004; Critchley & Garfinkel, 2017), ours is the first extensive approach to examine metabolic interoception in detecting glucose changes with EMA and CGM. Additionally, interoceptive deficits are linked to higher BMI (Robinson et al., 2021), highlighting a connection between metabolic sensing and weight regulation (Khalsa et al., 2022; Petzschner et al., 2021) and point to a potential sex-specific risk factor for future studies in women, who have a higher prevalence of (comorbid) eating and mood disorders (Valente et al., 2017). In contrast, glucose metabolism did not moderate mood regulation in metabolically healthy individuals, as they were unrelated to insulin sensitivity or BMI. We conclude that interoceptive accuracy helps regulate mood by enhancing responsiveness to metabolic signals circumstances (Füstös et al., 2013).

Despite showing that metabolic state ratings fully account for mood-related effects of hunger, limitations remain. First, we focus on glucose is one signal related to energy metabolism while additional endocrine signals, such as ghrelin (Fahed et al., 2023) or sex hormones (Soares et al., 2002; Steiner, 2003) also play a role. Future research may examine additional physiological markers, such as continuous monitoring of cortisol (Kusov et al., 2023), to expand our work. Second, while we found an association between glucose levels and mood in healthy participants, it remains unclear whether such associations would arise in individuals with metabolic or mental disorders (Ali et al., 2006; Anderson et al., 2001; Jones et al., 2021). Future research in clinical samples will help clarify a potential role of interoception in aberrant mood regulation.

To summarize, metabolic state ratings mediate the link between glucose levels and mood. Using a highly powered deep phenotyping design with repeated CGM and self-reported metabolic and mood measures in naturalistic settings, we further show that higher interoceptive accuracy is associated with lower mood fluctuations, highlighting the role of metabolic interoception in mood regulation. Our findings suggest that conscious signals of the body’s metabolic state affect mood in metabolically healthy participants, aligning with theories where changes in mood drive behavioral adjustments. This advances our understanding of mood regulation and supports the psychological construction of emotion theory, emphasizing interoception’s role in maintaining allostasis.

## Acknowledgment

We thank Antonia Schlaich, Johanna Voß, Franziska Peglow, Hannah Groß, Sophie Mathis, David Bartsch, Rauda Fahed, Katharina Bertl, Johanna Theuer, Ebru Sarmisak, Laura Heidiri, and Ricarda Bode for help with data acquisition and Alica Guzman for support in preprocessing the glucose data. The study was supported by the German Research Foundation (DFG) grants KR 4555/7-1, KR 4555/9-1, KR 4555/10-1 and DE 2319/22-1.

## Author contributions

NBK was responsible for the study concept and design. MK, MG, KK, & AK collected data under supervision by NBK and BD. AK, MG, BD, & NBK conceived the method and MBT and AK preprocessed the data. MK, KK, & NBK performed the data analysis. KK, MK, & NBK wrote the manuscript. All authors contributed to the interpretation of findings, provided critical revision of the manuscript for important intellectual content, and approved the final version for publication.

