## SupplementaryMaterial for "Glucose levels are associated with mood, but the association is mediated by ratings of metabolic state"

Kristin Kaduk^1^, Alessandro Petrella^1^, Sophie J. Müller^1^, Julian Koenig^2^, &

Nils B. Kroemer^1,3,4*^

Kristin Kaduk^1 #^, Marie Kaeber^1,#^, Anne Kühnel^2^, María Berjano Torrado^1^, Melina Grahlow^1^, Birgit Derntl^1,3^, & Nils B. Kroemer^1-3*^

^#^	equal contribution

^1^	Department of Psychiatry and Psychotherapy, Tübingen Center for Mental Health, University of Tübingen, Tübingen, Germany

^2^	Section of Medical Psychology, Department of Psychiatry and Psychotherapy, Faculty of Medicine, University of Bonn, Bonn, Germany

^3^	German Center for Mental Health (DZPG), partner site Tübingen

### Supplementary Material A: visualization of the Generalized Additive model (GAM) (related to Figure 3)

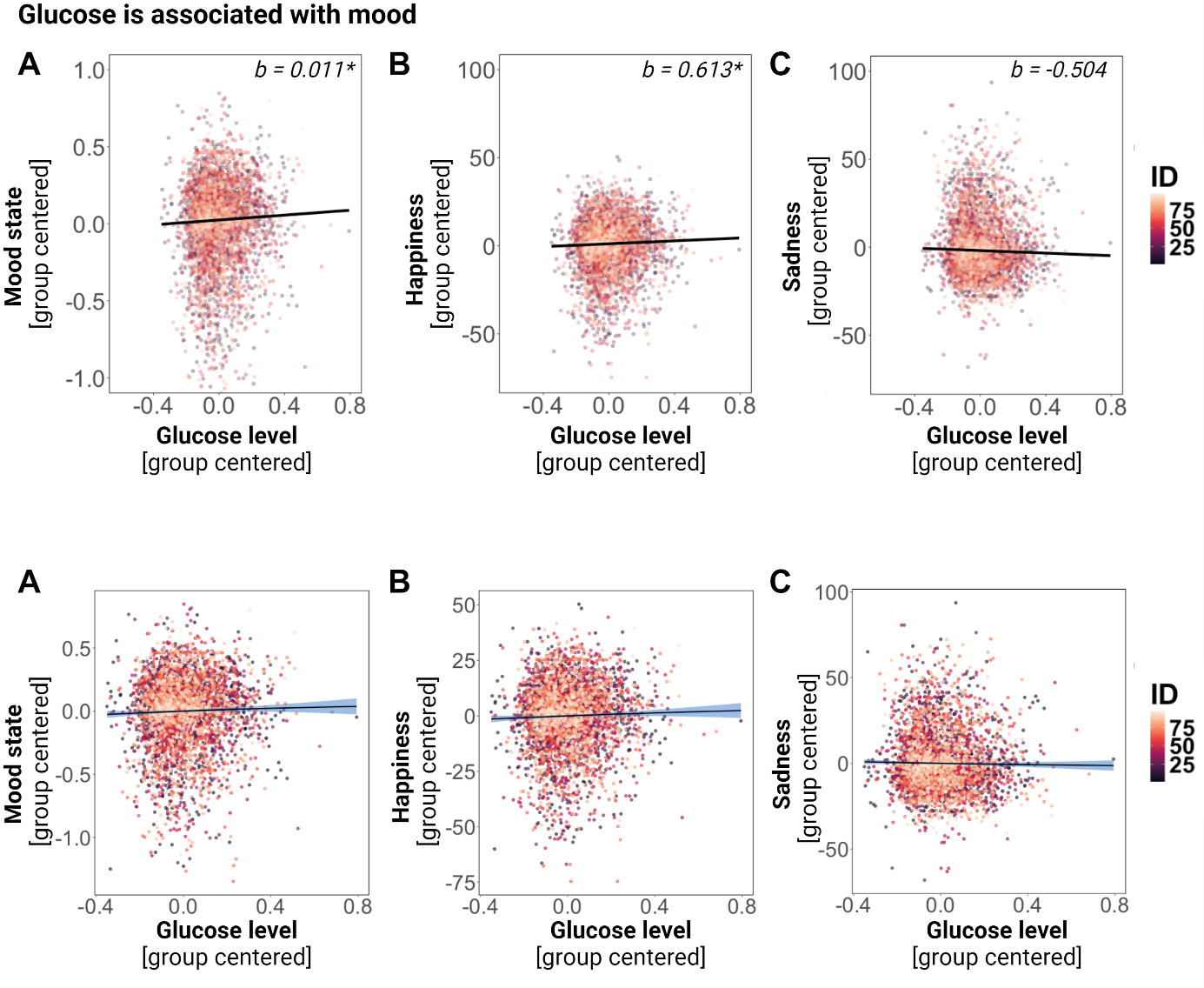

**Figure S1. Glucose levels are associated with mood, independent of metabolic state.** The graphs display the relationship between glucose levels and **A:** mood state, **B:** happiness, and **C:** sadness, indicating that when glucose levels are high, participants reported higher mood. Blue lines indicate the relationship (fixed effect) across participants estimated by a Generalized Additive Model (GAM). We used a Generalized Additive Model (GAM) visualization to examine the non-linearity of the data, which confirmed that the relationship between glucose and mood could be adequately described by a linear model. Each dot depicts an observation with a color-coding of participants. While mood state, happiness, and sadness are group centered on ID for visualization purposes, glucose level is log-transformed and group centered.
